# Novel lentiviral constructs for the specification of rare and ectopic cell types in human colonoids

**DOI:** 10.1101/2025.10.16.680258

**Authors:** Pritiprasanna Maity, Na Qu, Jorge O. Múnera

**Affiliations:** Department of Regenerative Medicine and Cell Biology, Medical University of South Carolina, Charleston, SC 29425

**Keywords:** human colonoids, tuft cells, goblet cells, M-cells, enteroendocrine cells

## Abstract

Human colonoids derived from adult intestinal tissue provide powerful models for epithelial biology but lack several rare or regionally restricted cell types. To overcome this limitation, we engineered doxycycline-inducible lentiviral constructs to drive expression of transcription factors that control lineage specification, including KLF4, POU2F3, SPIB, NEUROG3, and NEUROG3 T2A NKX6-3. Transduction of human colonoids with these constructs revealed that transient KLF4 induction is sufficient to generate TFF3 and WFDC2 expressing goblet cells. Unexpectedly KLF4 also induced GUCA2A positive colonocytes. In contrast, SPIB and POU2F3 were not sufficient to specify M-cells or mature tuft cells respectively, suggesting the requirement for additional permissive cues. Notably, co-expression of NEUROG3 and NKX6-3 induced enteroendocrine subtypes characteristic of the duodenum. These findings establish a set of lentiviral tools for induction of specific epithelial lineages in human colonoids providing a platform for replenishing cell types that are lost or altered in intestinal diseases.

## Introduction

In the past 15 years, organoid models have revolutionized the study of the human gastrointestinal (GI) epithelium[1]. Organoids can be derived from patient tissue and grown as strictly epithelial structures referred to as enteroids and colonoids [2]. Despite their utility, these models often lack rare epithelial cell types. We hypothesized that induced expression of key transcription factors (TFs) would be sufficient to specify a subset of these missing lineages.

To test this, we engineered lentiviral constructs allowing doxycycline-inducible expression of TFs known to regulate lineage specification. Human colonoids were transduced with these constructs, and their capacity to induce distinct differentiated cell types was assessed. Specifically, we cloned cDNAs for transcription factors implicated in cell fate determination: KLF4 which is required for goblet cell differentiation[4], SPIB which is required for M-cell differentiation[5], and POU2F3 which is required for tuft cell differentiation[6], into a doxycycline inducible lentiviral vector. For KLF4 we used isoform 2 (shorter isoform) cDNA. A previously validated NEUROG3 lentiviral construct, which induces enteroendocrine cells (EECs) in pluripotent stem cell-derived GI organoids[7], was also employed. Since NEUROG3 is required for general EEC specification in the GI tract[8], we further designed a NEUROG3-T2A-NKX6-3 construct to test whether co-expression with NKX6-3—a factor required for generating gastrin-producing G cells and somatostatin-producing D cells[9]—could direct differentiation toward duodenal EEC subtypes.

## Results

All constructs were packaged into lentiviral particles and used to transduce human enteroids and colonoids. To evaluate expression efficiency and determine optimal pulse-chase timing for differentiation, we applied a 24-hour doxycycline pulse followed by varied chase periods (Supplementary Figure 1A). Immediately after pulsing, we observed induction of the transgenes in 68% (KLF4), 65% (SPIB), 36% (NEUROG3), 34% (NEUROG3-T2A-NKX6-3), and 30% (POU2F3) of cells (Supplementary Figure 1B–K). Following doxycycline withdrawal, TF expression declined markedly, indicating tight control and minimal leakiness of the lentiviral system.

We next performed bulk RNA sequencing to define the global transcriptional effects of each TF. Principal component analysis (PCA) showed that doxycycline-treated colonoids separated from vehicle-treated controls along PC1, which accounted for 92% of variance (Supplementary Figure 2A). Moreover, within the doxycycline-treated group, samples clustered according to the specific TF expressed (Supplementary Figure 2B), confirming transcriptional divergence between induced colonoids.

KLF4 induction led to upregulation of goblet cell markers (Supplementary Figure 2C). Surprisingly, however, *MUC2* expression was not significantly increased, suggesting that KLF4 drives goblet cell subtypes distinct from canonical MUC2+ cells. To assess whether vehicle-treated colonoids were otherwise differentiated, we examined expression of colonocyte markers, including both general and pH-sensing subtypes. Canonical colonocyte markers *CA1, CA2*, and *DPP4* were highly expressed, consistent with terminal differentiation (Supplementary Figure 2D). Unexpectedly, KLF4 induction also led to robust upregulation of pH-sensing colonocyte genes, including *GUCA2A, GUCA2B, OTOP2*, and *OTOP3*.

Given recent findings that SPIB can promote pH-sensing colonocyte differentiation[10], we compared normalized RNA-seq counts between KLF4-induced ileal enteroids and SPIB-induced colonoids (Supplementary Figure 2E–F). Strikingly, *GUCA2A* and *GUCA2B* mRNA levels in KLF4-induced colonoids reached levels comparable to those in SPIB-induced conditions, revealing an unexpected role for KLF4 in inducing this lineage.

Together, these results indicate that KLF4 is sufficient to activate transcriptional programs associated with both mucus-secreting and pH-sensing colonocytes (Figure 1A). To confirm goblet cell induction, we performed immunostaining for TFF3 and WFDC2 following a 1-day doxycycline pulse and 4–6 day chase. KLF4-induced colonoids contained abundant TFF3+ and WFDC2+ goblet cells (Figure 1B–E). To validate GUCA2A+ colonocyte induction, we performed immunostaining and observed GUCA2A+ cells in KLF4-induced colonoids (Figure 1F–G), at levels comparable to those seen in SPIB-induced cultures (Figure 1H–I). These findings demonstrate that KLF4 promotes both goblet cell subtype differentiation and GUCA2A+ colonocyte identity.

**Figure 1.**
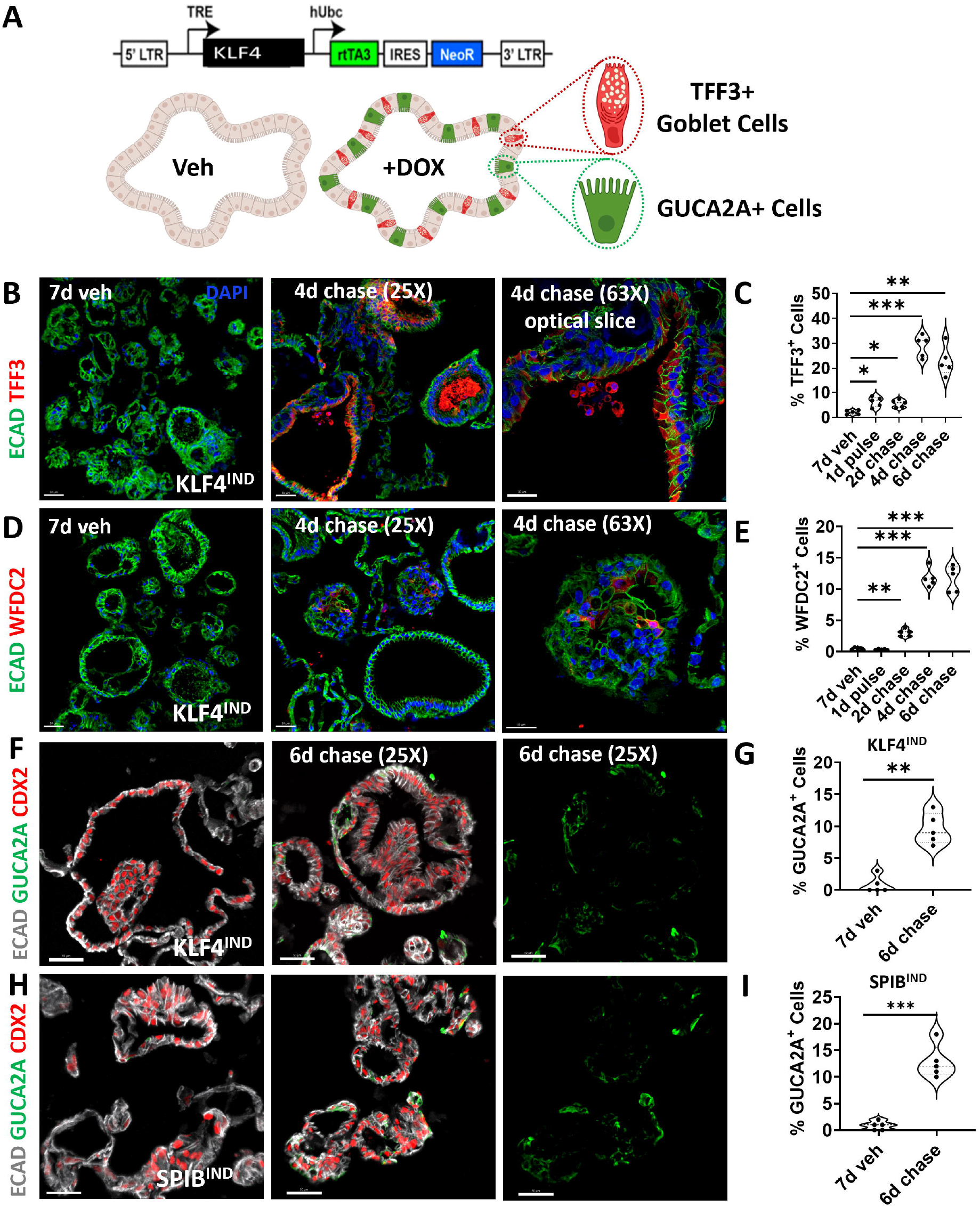
Transient KLF4 expression induces goblet cells subtypes and GUCA2A expressing colonocytes. (A) Schematic of KLF4 inducible (KLF4^IND^) construct. (B-E) TFF3 and WFDC2 staining in KLF4^IND^ colonoids. Staining for CDX2 and GUCA2A in (F-G) KLF4^IND^ and (H-I) SPIB^IND^ colonoids. Significance in (C,E) was determined by one-way ANOVA analysis with Tukey’s multiple comparisons test with *p<0.05, **p<0.01, and ***p<0.001 (n=5 different experiments per condition). Significance in (G,I) was determined by t-test with **p<0.01 and ***p<0.001 (n=5 different experiments per condition). Scale bars: 50 µm in left and middle panels in (B,D,F,H), and 30 µm in the right (63X) panels in (B,D). Schematic in (A) was generated using Biorender.

We next examined enteroendocrine cell differentiation NEUROG3-T2A-NKX6-3 induced colonoids (Supplementary figures 1G). We not only observed induction of pan-EEC mRNAs (*CHGA, CHGB)*, but we also observed expression of multiple small intestinal EEC markers (*GIP, SCT, NTS*), including duodenal-specific markers (*MLN, GAST, SST, GHRL*) transcripts. We confirmed by immunostaining that CHGA, GAST, GIP, CCK and SST expressing cells were significantly increased by induced NEUROG3-T2A-NKX6-3 expression compared to induced NEUROG3 expression alone (Figure 2A-J) confirming that multiple ectopic duodenal EEC subtypes were induced in NEUROG3-T2A-NKX6-3^IND^ colonoids.

**Figure 2.**
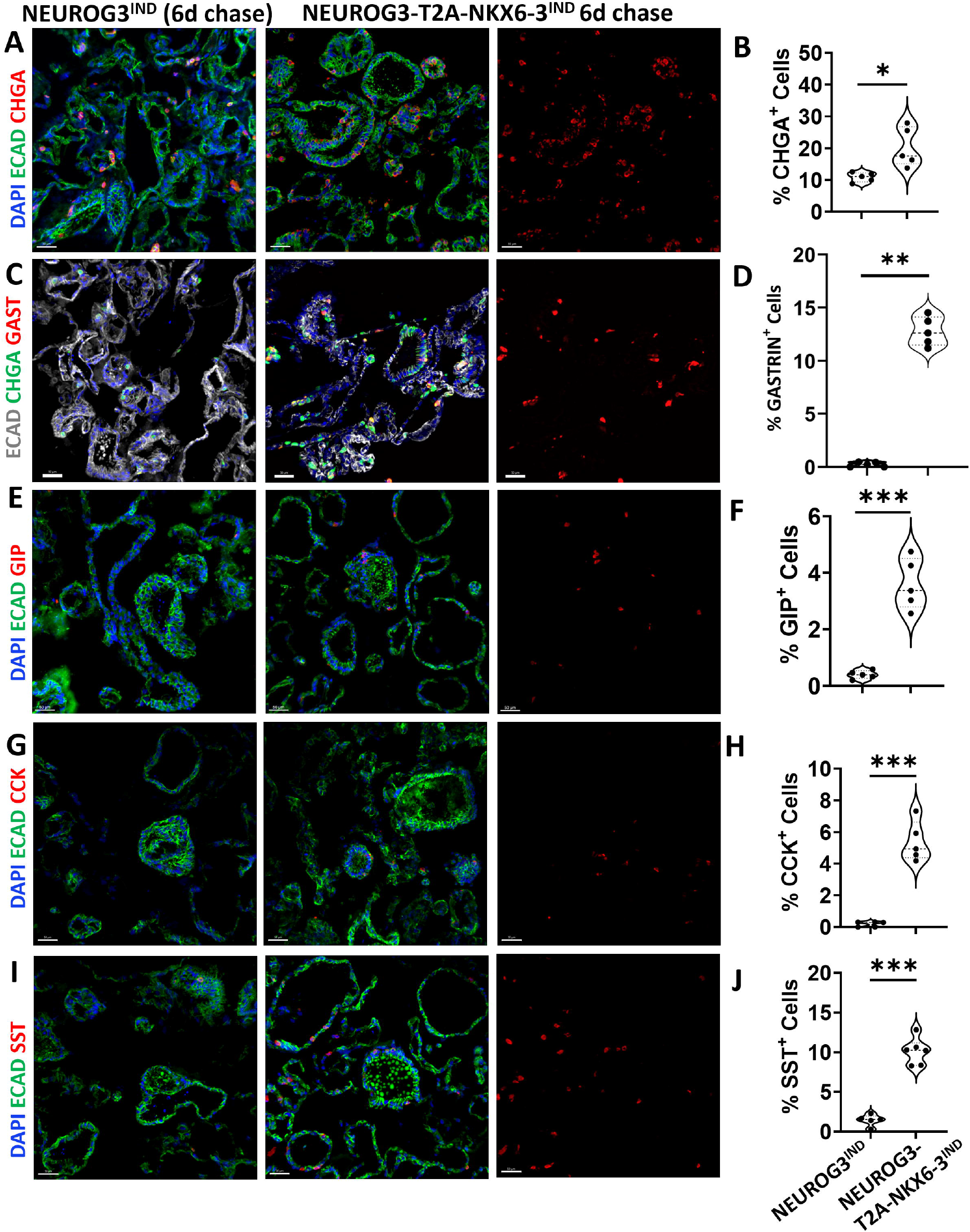
Generation of small intestinal and duodenal cell types in NEUROG3-T2A-NKX6-3^IND^ colonoids. Immunofluorescence staining and quantification for (A-B) CHGA, (C-D) GIP (E-F) CCK, and (G-H) SST in NEUROG3^IND^ and NEUROG3-T2A-NKX6-3^IND^ colonoids. Significance was determined by t-test with *p<0.05 and ***p<0.001 (n=5 different experiments per condition). Scale bars: 50 µm in all panels.

In contrast, we did not observe GP2+ M-cells at any time point in either colonoids (Supplementary figure 2A-C) or ileal enteroids (Supplementary figure 3F,G), despite a 65% and 50% induction of SPIB following a 1-day pulse respectively (Supplementary figures 1D,E, and supplementary figure 3D-E). Similarly, POU2F3 induction (Supplementary Figure 4A) failed to elicit expression of tuft cell-specific mRNAs (Supplementary Figure 4B). Unexpectedly, we observed pEGFR+ cells in POU2F3-induced colonoids (Supplementary Figures 4C–E), suggesting that POU2F3 is not sufficient to generate fully mature tuft cells under these conditions.

## Discussion

In conclusion, we have developed lentiviral tools that generate human colonoids that are enriched for differentiated cell types including ectopic duodenal EECs. The KLF4^IND^ construct will allow the interrogation of regional or crypt-base versus crypt-top differences in goblet cells as well as KLF4 mediated induction of GUCA2A+ colonocytes. The failure of our POU2F3^IND^ and SPIB^IND^ constructs to generate tuft cells and M-cells respectively, suggests that either permissive signals and/or other TFs may be necessary to induce these cell types, consistent with recent reports[10, 11]. Our NEUROG3-T2A-NKX6-3^IND^ construct indicates that by utilizing the correct combination of TFs, we can generate EEC subtypes of interest. This work will inform strategies for the replenishment of cell types that are depleted in conditions such as inflammatory bowel diseases.

## References

[1] Kretzschmar K, Clevers H. Organoids: Modeling Development and the Stem Cell Niche in a Dish. Dev Cell 2016;38(6):590–600.

[2] Stelzner M, Helmrath M, Dunn JC, Henning SJ, Houchen CW, Kuo C, Lynch J, Li L, Magness ST, Martin MG, Wong MH, Yu J, Consortium NIHISC. A nomenclature for intestinal in vitro cultures. Am J Physiol Gastrointest Liver Physiol 2012;302(12):G1359–63.

[3] Beumer J, Puschhof J, Bauza-Martinez J, Martinez-Silgado A, Elmentaite R, James KR, Ross A, Hendriks D, Artegiani B, Busslinger GA, Ponsioen B, Andersson-Rolf A, Saftien A, Boot C, Kretzschmar K, Geurts MH, Bar-Ephraim YE, Pleguezuelos-Manzano C, Post Y, Begthel H, van der Linden F, Lopez-Iglesias C, van de Wetering WJ, van der Linden R, Peters PJ, Heck AJR, Goedhart J, Snippert H, Zilbauer M, Teichmann SA, Wu W, Clevers H. High-Resolution mRNA and Secretome Atlas of Human Enteroendocrine Cells. Cell 2020.

[4] Katz JP, Perreault N, Goldstein BG, Lee CS, Labosky PA, Yang VW, Kaestner KH. The zinc-finger transcription factor Klf4 is required for terminal differentiation of goblet cells in the colon. Development 2002;129(11):2619–28.

[5] Kanaya T, Hase K, Takahashi D, Fukuda S, Hoshino K, Sasaki I, Hemmi H, Knoop KA, Kumar N, Sato M, Katsuno T, Yokosuka O, Toyooka K, Nakai K, Sakamoto A, Kitahara Y, Jinnohara T, McSorley SJ, Kaisho T, Williams IR, Ohno H. The Ets transcription factor Spi-B is essential for the differentiation of intestinal microfold cells. Nat Immunol 2012;13(8):729–36.

[6] Gerbe F, Sidot E, Smyth DJ, Ohmoto M, Matsumoto I, Dardalhon V, Cesses P, Garnier L, Pouzolles M, Brulin B, Bruschi M, Harcus Y, Zimmermann VS, Taylor N, Maizels RM, Jay P. Intestinal epithelial tuft cells initiate type 2 mucosal immunity to helminth parasites. Nature 2016;529(7585):226–30.

[7] McCracken KW, Cata EM, Crawford CM, Sinagoga KL, Schumacher M, Rockich BE, Tsai YH, Mayhew CN, Spence JR, Zavros Y, Wells JM. Modelling human development and disease in pluripotent stem-cell-derived gastric organoids. Nature 2014;516(7531):400–4.

[8] Jenny M, Uhl C, Roche C, Duluc I, Guillermin V, Guillemot F, Jensen J, Kedinger M, Gradwohl G. Neurogenin3 is differentially required for endocrine cell fate specification in the intestinal and gastric epithelium. EMBO J 2002;21(23):6338–47.

[9] Choi MY, Romer AI, Wang Y, Wu MP, Ito S, Leiter AB, Shivdasani RA. Requirement of the tissue-restricted homeodomain transcription factor Nkx6.3 in differentiation of gastrin-producing G cells in the stomach antrum. Mol Cell Biol 2008;28(10):3208–18.

[10] Wang D, Spoelstra WK, Lin L, Akkerman N, Krueger D, Dayton T, van Zon JS, Tans SJ, van Es JH, Clevers H. Interferon-responsive intestinal BEST4/CA7(+) cells are targets of bacterial diarrheal toxins. Cell Stem Cell 2025;32(4):598–612 e5.

[11] Huang L, Bernink JH, Giladi A, Krueger D, van Son GJF, Geurts MH, Busslinger G, Lin L, Begthel H, Zandvliet M, Buskens CJ, Bemelman WA, Lopez-Iglesias C, Peters PJ, Clevers H. Tuft cells act as regenerative stem cells in the human intestine. Nature 2024;634(8035):929–35.

