## Supplementary information for "Novel lentiviral constructs for the specification of rare and ectopic cell types in human colonoids"

### SUPPLEMENTARY FIGURE LEGENDS

**Supplementary figure 1. Induction efficiencies of transcription factors in human colonoids.** (A) Schematic of pulse-chase experiments**.** Immunostaining results and quantification from a time course of induced (B,C) POU2F3, (D,E) SPIB, (F,G) KLF4, (H,I) NEUROG3 and (J,K) NEUROG3-T2A-NKX6-3 in colonoids. Significance was determined by one-way ANOVA analysis with Tukey's multiple comparisons test with *p<0.05, **p<0.01, ***p<0.001 and ****p<0.0001 (n=3-5 different experiments per condition). Scale bars: 50 μm in all image panels.

**Supplementary figure 2. Transcriptomic analysis confirms the specification of some targeted cell types.** (A) Principal component analysis (PCA) of RNA seq data from vehicle controls and DOX induced colonoids. (B) PCA analysis of DOX induced colonoids. Heatmaps generated from transcript counts from RNA-seq data in control and DOX induced colonoids for (C) goblet cell, (D) colonocyte subtypes and (G) enteroendocrine cell mRNAs. (E,F) Graphs generated from transcript counts for *GUCA2A* and *GUCA2B* from RNA-seq data from KLF4 induced colonoids and SPIB induced ileal enteroids, Significance was determined by one-way ANOVA analysis with Tukey's multiple comparisons test with *p<0.05, **p<0.01, ***p<0.001 and ****p<0.0001.

**Supplementary figure 3. SPIB is not sufficient for the induction of M-cells in colonoids or ileal enteroids.** (A) Schematic of generation of $SPIB^{IND}$ colonoids and ileal enteroids. (B,C) Immunostaining and quantification of the M-cell marker GP2 in $SPIB^{IND}$ colonoids. (D, E) Immunostaining and quantification of SPIB in a pulse-chase time course of $SPIB^{IND}$ ileal enteroids. (F,G) Immunostaining and quantification of GP2 in a pulse-chase time course of $SPIB^{IND}$ ileal enteroids. Significance was determined by one-way ANOVA analysis with Tukey's multiple comparisons test with ***p<0.001 (n=5 different experiments per condition). Scale bars: 50 μm in image panels. Schematic in (A) was generated using Biorender.

**Supplementary figure 4. Induction of POU2F3 is not sufficient for the differentiation of mature tuft cells.** A) Schematic of POU2F3 inducible ($POU2F3^{IND}$) construct. (B) Heatmap generated from transcript counts from RNA-seq data, of tuft cell genes in control and DOX induced colonoids. Immunostaining and quantification for (C-E) Phospho-EGFR in $POU2F3^{IND}$ colonoids. Significance was determined by one-way ANOVA analysis with Tukey's multiple comparisons test with *p<0.05, **p<0.01, ***p<0.001 and ****p<0.0001 (n=3-5 different experiments per condition). Scale bars: 50 μm in image panels in (C), 30 μm in (D).

### METHODS

#### Generation of lentiviral constructs.

The pINDUCER20 (Plasmid #44012) was purchased from Addgene. The cDNA sequences for POU2F3, SPIB, KLF4 (NM_004235.5), NEUROG3 and NEUROG3-T2A-NKX6-3 were synthesized by Geneart (Thermo fisher Scientific) and cloned into pDONOR221. The cDNAs were then cloned into pINDUCER20 through Gateway™ cloning using the manufacturer's protocol. The resulting pINDUCER20 constructs were then re-sequenced. Plasmids were isolated using the Qiagen HiSpeed® Plasmid Midi Kit.

**Generation and culturing of colonoids and enteroids .**

Human healthy ileal enteroids, along with healthy colonoids, were obtained from Mindy Engevik and the Baylor College of Medicine 3D organoids Core respectively. These enteroids and colonoids were cultured as previously described [1]. Enteroids and colonoids were split once a week by enzymatic (TrypLE) dissociation. L-WRN condition media (L-WRN cells, ATCC CRL-3276TM) was prepared as per the ATCC protocol. The enteroids and colonoids were grown in expansion media composed of Advanced DMEM/F12 (Gibco) and L-WRN condition media (CM) (1:1 ratio) with 10 μM Y27632 (Tocris Cat# 1254), 10 μM SB431542 (Tocris Cat# 1614/50), 1mM N-acetyl cysteine (Sigma-Aldrich Cat# A9165), 50 ng/mL EGF (R&D systems), 10 mM nicotinamide (Sigma-Aldrich Cat# N0636), 500 nM A83-01 (Tocris Cat# 2939) and 10 μM SB202190 (Sigma-Aldrich Cat# S7067) to support the growth and long-term cultures of 3D colonoids and ileal enteroids.

**Lentivirus packaging and infection of colonoids and ileal enteroids.**

For packaging of lentivirus, 293T cells were purchased from ATCC. 293T cells cultured in 60 mm plates until they reached 60-70% confluency. On day 1, pMD2.G (Addgene #12259), pSPAX2 (Addgene #12260)and target DNA (KLF4, POU2F3, SPIB, NEUROG3 and NEUROG3-T2A-NKX6-3) were suspended in 350 μl Opti-MEM media (Gibco 11058) and for control, only plasmid DNA was suspended in the Opti-MEM media. PEI was then added and incubate for 15 minutes. The mixture was then added drop by drop around the tissue culture dish and mixed by gently swirling the plate. On the day 2 the media was replaced with colonoids and enteroids expansion media without antibiotics and cultured for an addditional 24 hrs. On day 3 , media containing virus particles was collected and filtered through a 0.45μm membrane. Polybrene was added (8ug/ml) to the filtered viral media to enhance the infection.

For lentiviral infection, human enteroids and colonoids were cultured in 24-well plates, with approximately 100 organoids per well. Two wells were dissociated into single cells using TrypLE Express (Life Technologies) supplemented with 10 μM ROCK inhibitor (Y-27632, Tocris Cat# 1254) and incubated at 37°C for 5 minutes. Mechanical trituration was performed by pipetting to ensure complete dissociation. The reaction was quenched by adding an equal volume of culture medium, and cells were pelleted at 300 × g for 3 minutes. Approximately 60,000–100,000 single cells were resuspended in a 1:1 mixture of enteroid expansion media and viral supernatant and transferred to ultra-low attachment 6-well plates (Corning, Cat# 3471). The cells were incubated with virus at 37°C for 6 hours. Following infection, the suspension was centrifuged and embedded in Matrigel for 3D culture. After 4 days in expansion media, G418 (63 μg/mL) was added for selection over 5 days. Organoids were then maintained in antibiotic-free expansion media. Infected enteroids and colonoids were used for one to two experimental replicates before being discarded. For subsequent experiments, fresh organoid cultures from previously uninfected enteroids or colonoids were infected following the same procedure.

**Induction of targeted cell types in human colonoids and enteroids.**
Colonoids and ileal enteroids were enzymatically dissociated into single cells. The cell suspension was mixed with Matrigel and seeded into the 24 well plate (15 µl/ well) with approximately 5,000 cells plated per well. Cells were then fed with expansion media. After 4 days, the colonoids and ileal enteroids were pulsed with DMSO (vehicle) or 1 µg/ml Doxycycline (DOX) for 24 hrs in expansion media which was then replaced with differentiation media containing advanced DMEM/F12, 3% conditioned media, 10 µM Y27632 and 1mM NAC. The colonoids and ileal enteroids were collected directly after the pulse and after 2-, 4- and 6-days post-pulse for downstream analyses.

**RNA isolation and Bulk RNA-seq sequence assembly abundance estimation**

RNA was extracted using NucleoSpin® RNA extraction kit (Macharey-Nagel) according to manufacturer's protocols. RNA library construction and RNA sequencing was performed by Innomics Inc using the DNBseq platform. QC analysis was performed with data from the DNBseq platform, adapter sequences were removed and data was provided as FASTQ files. Downstream analysis was performed using Partek® Flow® software. Heatmaps were generated using TPM tables that were then converted into heatmaps using Morpheus (Broad Institute). The gene lists for heatmaps were derived based on a previous study [2].

**Immunostaining and Quantification.**

Colonoids and ileal enteroids were fixed in 4% paraformaldehyde (PFA) for 1 hour on ice. The PFA fixed colonoids and ileal enteroids were washed with cold PBS, resuspended in 30% sucrose and incubated overnight at 4°C on a rocking platform. The colonoids and ileal enteroids were then frozen in OCT. OCT sections were blocked using blocking solution (5% donkey serum in 1× PBS plus 0.5% Triton-X) for 30 min and incubated overnight at 4 °C with primary antibodies diluted in blocking solution. Slides were then washed 3X with 1X PBS plus 0.5% Triton-X and incubated with secondary antibodies with DAPI in blocking buffer for 2 hr at room temperature. Slides were then washed 3X with 1X PBS plus 0.5% Triton-X and mounted using Fluoromount-G® (SouthernBiotech, Birmingham, AL, USA). The primary antibodies used were goat anti-E-Cadherin (1:400, R&D, AF648), rabbit anti-POU2F3 (1:200, Sigma, HPA019652), rabbit anti-P-EGFR (1:100, Abcam, ab205828), rabbit anti-KLF4 (1:100, Sigma, HPA002926), rabbit anti-TFF3 (1:200, Sigma, HPA035464), rabbit anti-WFDC2 (1:100, Sigma, HPA042302), rabbit anti-GUCA2A ( 1:300, Sigma, HPA018215), mouse anti-CDX2 (1:300, Biogenex, MU392A5UC), sheep anti-NEUROG3 (1:100, R&D, AF3444-SP), rabbit anti-GASTRIN (1:100, Cell Marque, 256A-14), mouse anti-CHROMOGRANIN A (1:250, DSHB, CPTC-CHGA-2), rabbit anti-CCK (1:200, Sigma, HPA069515), rabbit anti-SOMATOSTATIN (1:100, Cell Marque, 332R-14), rabbit anti-5-HT (SEROTONIN) (1:100, Immunostar, 20080), rabbit anti-GIP (1:200, Sigma, HPA021612), rabbit anti-GLP1 (1:500, Abcam, ab108443), rabbit anti-SPIB (1:100, Cell Signaling, 14323s) and rabbit anti-GP2 (1:200, Sigma, HPA015739. The secondary antibodies were donkey anti-goat; Alexa Fluor 488 (1:500, Fisher, A11055), donkey anti-mouse; Alexa Fluor 568 (1:500, Fisher, A10037) and donkey anti-rabbit; Alexa Fluor 647 (1:500, Fisher, A31573). Imaging was performed on a Zeiss LSM 880 NLO with Airscan confocal microscope and analyzed using Imaris Imaging Software 9.9 (Oxford Instruments, Abingdon, UK). For each experimental condition, images were generated from minimum of 3 fields which were then averaged for each colonoid passage. Quantitative analysis was also performed on images ImageJ software The percentage of marker positive cells was calculated based on the total number of DAPI positive nuclei.

#### Statistical tests

All experiments with colonoids and ileal enteroids were repeated five times except for NEUROG3$^{IND}$ studies which were repeated 3 times. For each set of experimental conditions, 3-4 wells of colonoids or ileal enteroids were used. Quantification of data are represented as mean ± SEM unless otherwise specified. Significance was determined by either unpaired t tests with 2-tailed distribution and two-sample equal variance when comparing 2 conditions or one-way ANOVA with Tukey's multiple comparison test when comparisons >2 conditions. Significance is represented as $p<0.05$*,$<0.01$**,$p<0.001$*** and $p<0.0001$**** as denoted in Figure legends. Statistics were performed and graphs were generated in Graph Pad Prism. Some figure schematics were generated using BioRender.

### REFERENCES

[1] Sayed IM, Tindle C, Fonseca AG, Ghosh P, Das S. Functional assays with human patient-derived enteroid monolayers to assess the human gut barrier. *STAR Protocols.* 2. Elsevier Inc.; 2021:100680.

[2] Burclaff J, Bliton RJ, Breau KA, Ok MT, Gomez-Martinez I, Ranek JS, Bhatt AP, Purvis JE, Woosley JT, Magness ST. A Proximal-to-Distal Survey of Healthy Adult Human Small Intestine and Colon Epithelium by Single-Cell Transcriptomics. Cell Mol Gastroenterol Hepatol 2022;13(5):1554-89.

Supplementary figure 1. Induction efficiencies of transcription factors in human colonoids.

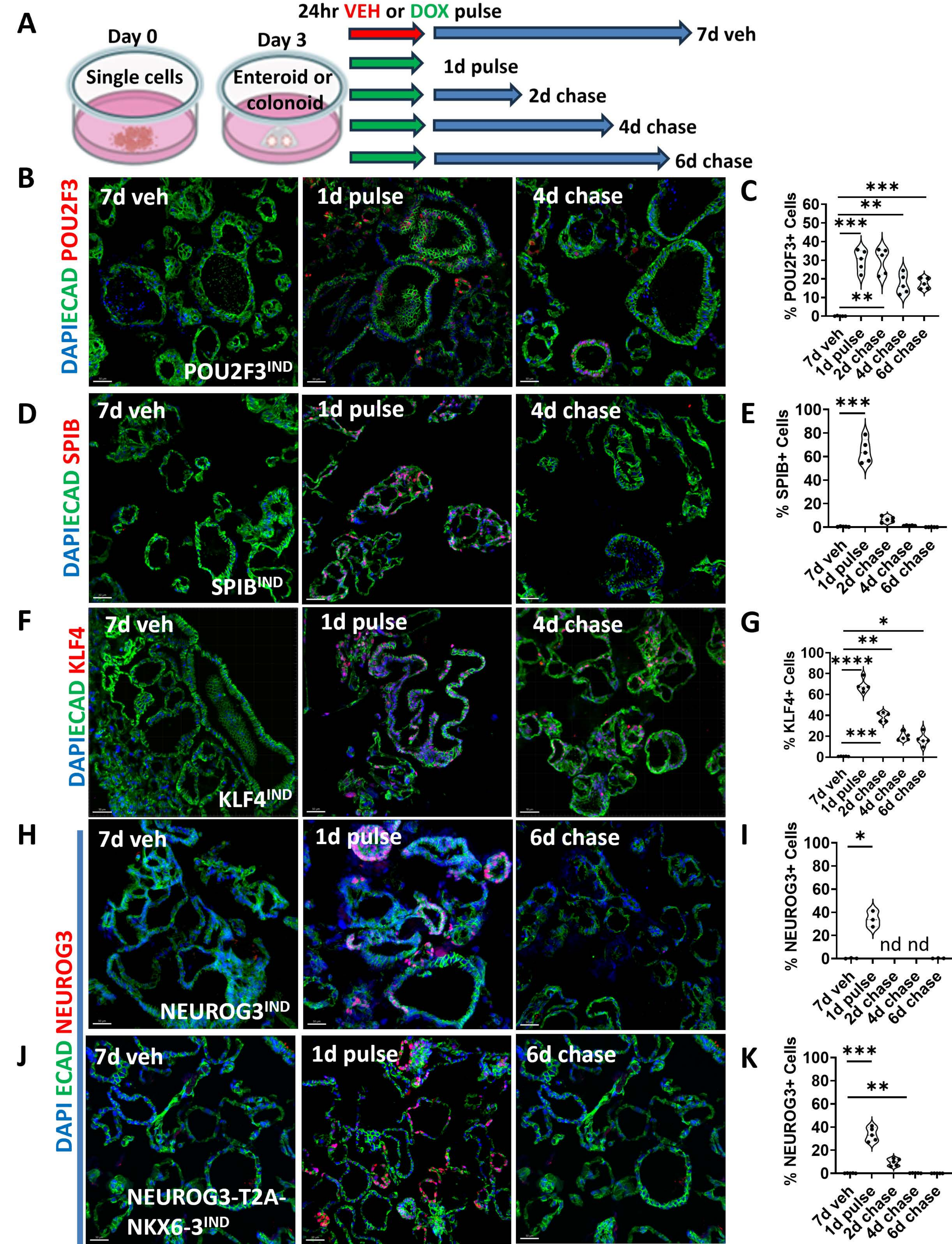

**Supplementary figure 2. Transcriptomic analysis of confirms the specification of some targeted cell types.**

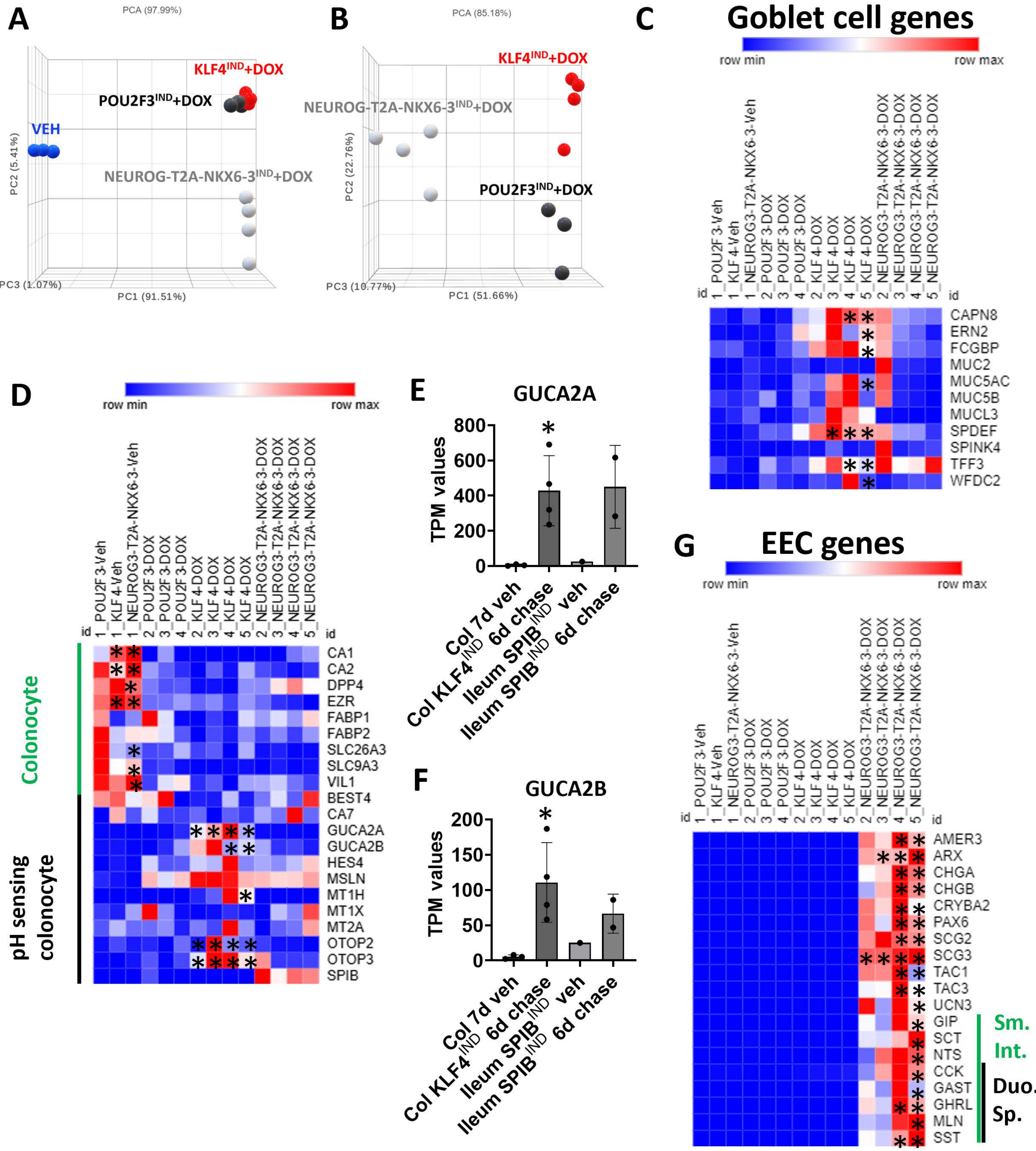

**Supplementary figure 3. SPIB is not sufficient for the induction M-cells in colonoids or ileal enteroids.**

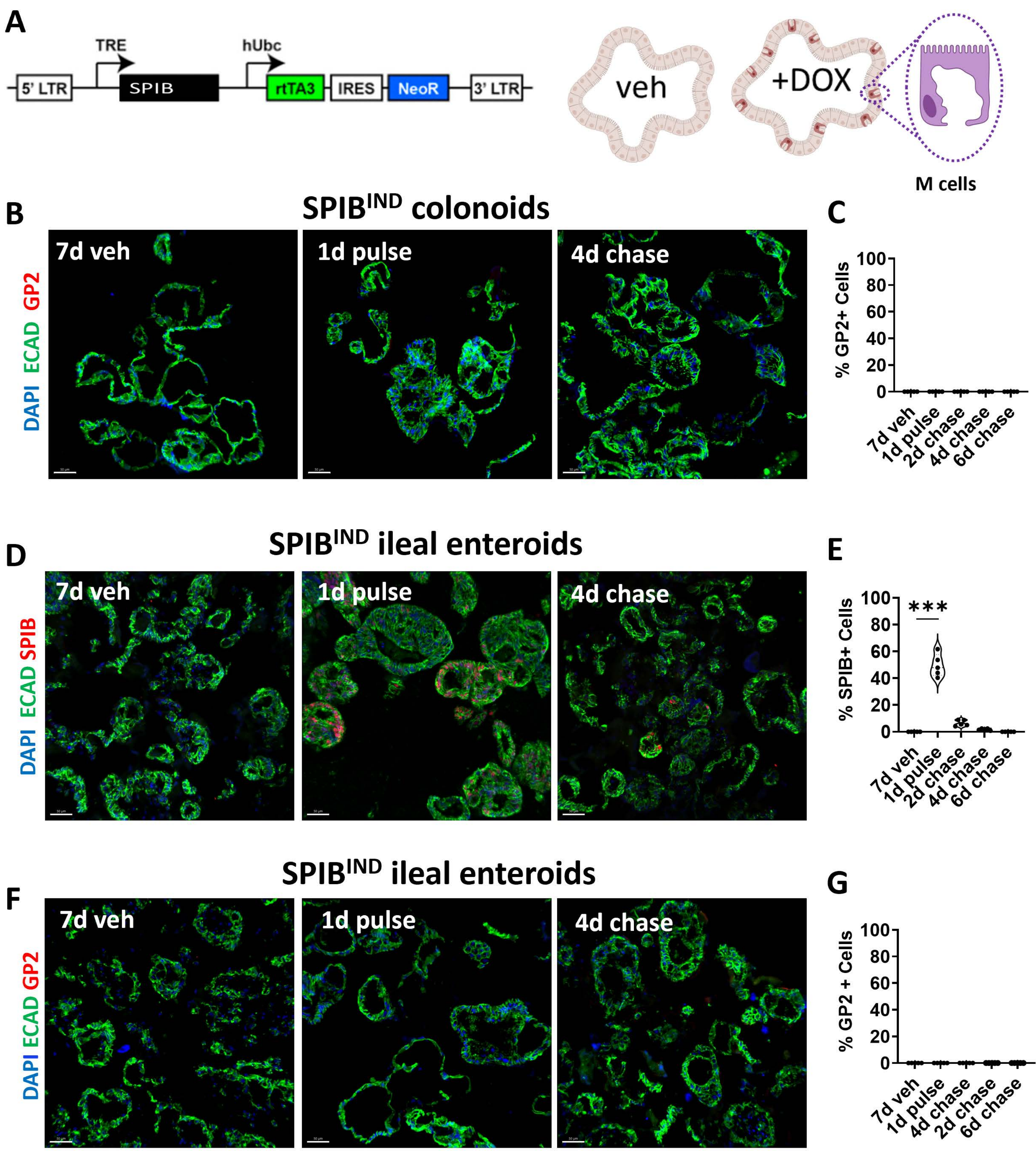

**Supplementary figure 4. POU2F3 is not sufficient for the induction tuft cells in colonoids.**

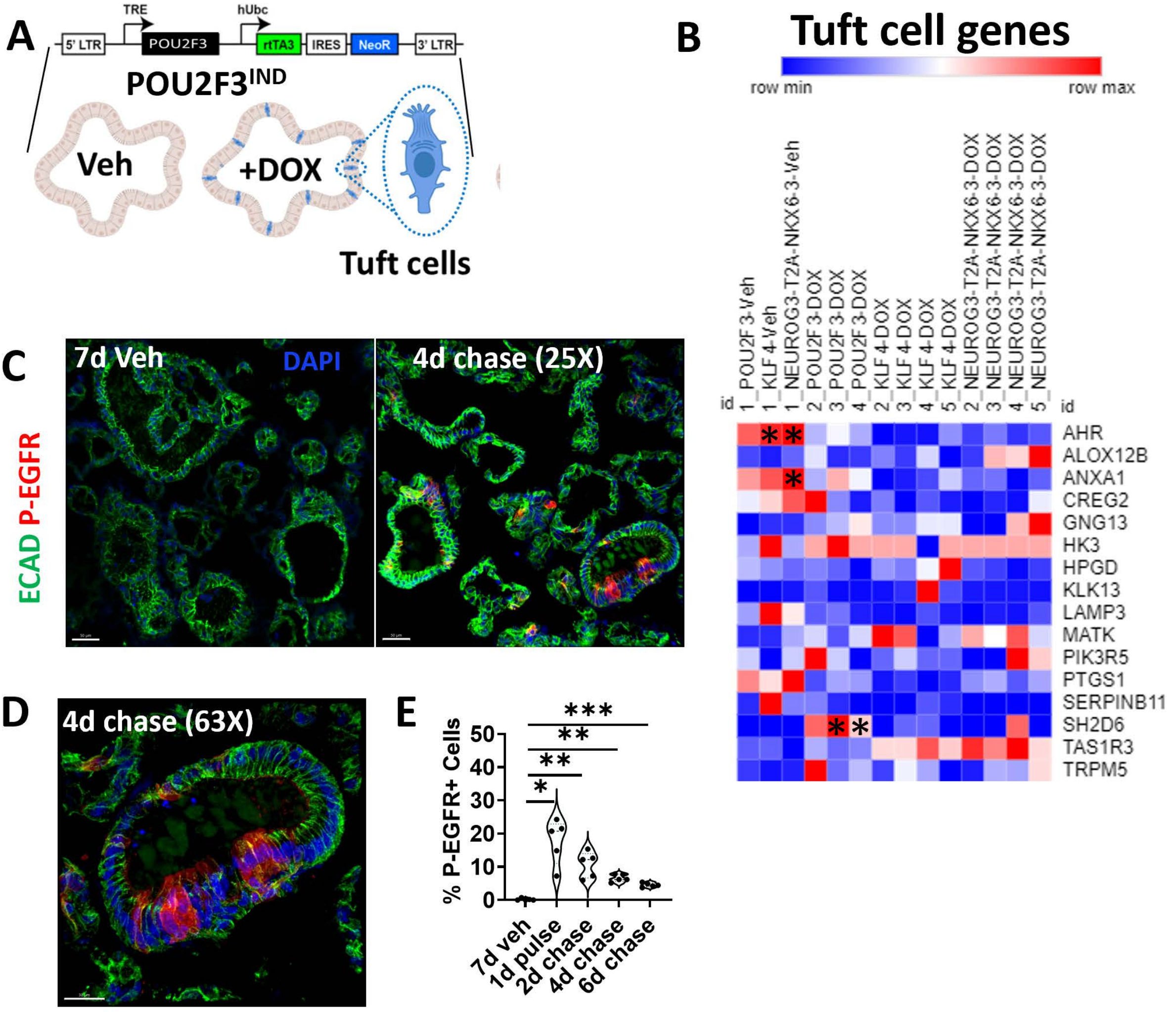
